# Rebuilding the autoimmune-damaged corneal stroma through topical lubrication

**DOI:** 10.1101/2024.11.29.626078

**Authors:** Yael Efraim, Feeling Yu Ting Chen, Seyyed Vahid Niknezhad, Dylan Pham, Ka Neng Cheong, Luye An, Hanan Sinada, Nancy A. McNamara, Sarah M. Knox

## Abstract

Corneal lubrication is the most common treatment for relieving the signs and symptoms of dry eye and is considered to be largely palliative with no regenerative functions. Here we challenge this notion by demonstrating that wetting the desiccated cornea of an aqueous-deficient mouse model with the simplest form of lubrication, a saline-based solution, is sufficient to rescue the severely disrupted collagen-rich architecture of the stroma, the largest corneal compartment that is essential to transparency and vision. At the single cell level we show that stromal keratocytes responsible for maintaining stromal integrity are converted from an inflammatory state into unique reparative cell states by lubrication alone, thus revealing the extensive plasticity of these cells and the regenerative function of lubricating the surface. We further show that the generation of a reparative phenotype is due, in part, to disruption of an IL1β autocrine amplification loop promoting chronic inflammation. Thus, our study uncovers the regenerative potential of topical lubrication in dry eye and represents a paradigm shift in our understanding of its therapeutic impact.

## Introduction

A healthy tear film provides an aqueous coating necessary for optimal vision and tissue function while also shielding the ocular surface from environmental, inflammatory, and microbial insult. Prolonged dryness, such as that resulting from aqueous-deficient dry eye, induces a vast array of pathological alterations in cell phenotypes and corneal structure. Despite the high prevalence of this chronic injury globally (1), there remains a glaring lack of effective treatment options. Consequently, the most common therapy used are over-the-counter topical aqueous lubricants aimed to relieve dry eye symptoms^1^. As the apical epithelial layers show very little to no recovery with routine application(2, 3), saline-based solutions are widely thought to have no regenerative properties. However, studies to date have overlooked the cellular compartment residing beneath the epithelium that is crucial to vision and comprises 90% of the corneal thickness - the collagen-rich stroma.

The stromal compartment plays a critical role in vision by maintaining corneal transparency, avascularity and refractive function, while providing the mechanical properties necessary for tissue strength and preservation of corneal shape(4, 5). These properties are primarily reliant on densely packed collagen fibrils (primarily types 1 and 5) arranged as lamellae parallel to the epithelial surface (6). These form a tight extracellular matrix (ECM) enriched in a diverse array of proteoglycans and glycoproteins, such as perlecan and thrombospondin, respectively, necessary for regulating the size and organization of the fibers as well as facilitating cell-to-cell and cell-to-matrix interactions (4). The primary cells involved in creating and maintaining the structural complexity of the stroma are the keratocytes. These are specialized, relatively quiescent neural-crest derived cells that, under homeostatic conditions, maintain the aligned nanofibrils and matrix composition. Keratocytes have been shown to exhibit lineage heterogeneity under healthy conditions, with single cell sequencing of murine and human stroma revealing 3 and 4 (7) distinct populations, respectively. Although no single cell analyses have been focused on the corneal stroma compartment to date, studies have demonstrated that in response to acute damage, keratocytes undergo apoptosis or transition into activated fibroblasts and myofibroblasts that possess reparative functions. This transitional phenotype is postulated to be attributed to stem cell-like properties derived from their cell of origin (8). However, the impact of chronic inflammatory disease, on the heterogeneity and plasticity of keratocytes, as well as the intricate signaling pathways governing their behavior, is unclear. Furthermore, despite their intimate relationship with the epithelium through the basement membrane, whether keratocytes transition into disruptive or regenerative cell states in response to changes at the ocular surface, such as those induced by simple lubrication, remains unknown.

Using the autoimmune-regulator knock out (Aire KO) mouse model to mimic many of the pathological features of dry eye (9), we have discovered that simply lubricating the ocular surface induces the generation of a heterogeneous pool of reparative keratocytes, transitioning them from inflammatory to reparative cells to drive stromal repair. Topical lubrication with preservative-free phosphate buffer saline (PBS), the simplest form of artificial tears commonly used by patients, restores stromal architecture to a homeostatic-like state. In addition, the basement membrane (BM) is also restored, suggestive of the re-establishment of matrix-cell communication. These outcomes are achieved, in part, by disrupting a chronically active IL1- MAPK signaling pathway that is among the most common mediators of tissue damage. Thus, contrary to our prevailing belief that topical saline-like lubricants for dry eye disease are largely palliative, we show that this treatment effectively reinstates key aspects of keratocyte function required for stromal regeneration.

## Results

### Lubrication restores the structural integrity of the stroma and the basement membrane

Reflex tear secretion in response to changes at the ocular surface is essential to maintain corneal homeostasis and promote wound healing, all of which are profoundly reduced in human patients with dry eye disease (10). Topical aqueous lubrication is a well-established palliative therapy for relieving symptoms of chronic dry eye (11) but is not an effective therapy to restore epithelial barrier function in moderate to severe cases (3). However, the therapeutic impact of topical lubrication on the epithelial BM and corneal stroma in dry eye is unknown.

To test whether lubrication possesses therapeutic properties in the corneal stroma, we utilized a well-characterized autoimmune-mediated dry eye mouse model deficient in the autoimmune regulator (*Aire*) gene that recapitulates many of the pathologies of patients with severe disease including chronic epithelial wound healing, reduced innervation, disrupted barrier function, and stromal scarring (9, 12, 13). *Aire* knockout (*KO*) female mice quickly develop dry eye over a 2- week period, with mildly reduced tear secretion and corneal barrier function at 5 weeks (wks) of age, followed by extensive exocrinopathy, and severe pathologies affecting the corneal epithelial cells, their basement membrane (BM), and the nerves that innervate them by 7 wks (9). Similar to severe human disease, we also found the stromal compartment to be severely disrupted over time (Figure 1B). In contrast to the narrow collagen fibers densely packed and uniformly aligned parallel to the corneal surface in the healthy tissue, desiccated tissue showed a significant increase in the frequency of thicker collagen fibers and in fiber angles relative to the epithelium (>40-90 degrees; Figure 1B, E and F and Supplementary Figure S1A-B). Strikingly, PBS restored stromal architecture, with collagen fiber assemblies resembling WT controls after 14 days of treatment (Figure 1B, E and F and Supplementary Figure S1A-B). Consistent with this rescue, stromal swelling (edema), an outcome associated with disruption of lamella and severe dry eye (14), was also rescued by treatment (Figure 1G). Finally, the nearly absent BM in the untreated controls, a structure required for numerous processes such as anchoring epithelial cells and maintaining barrier function (REF), was also restored, as shown by the renewed deposition of two essential BM proteins, perlecan and laminin, along the basal epithelium (Figure 1B, C, D).

**Figure 1:**
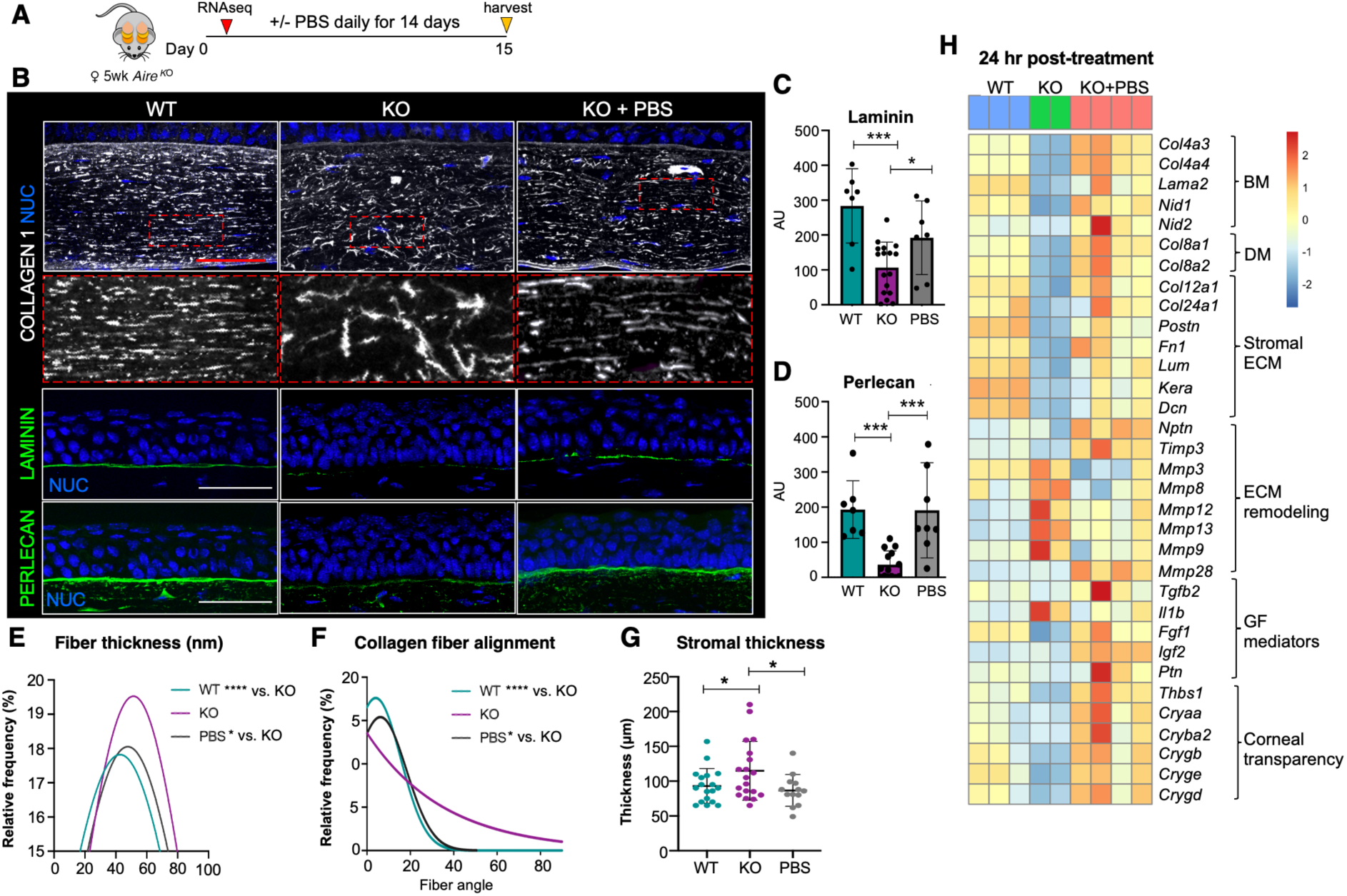
Topical lubrication of the ocular surface promotes stromal regeneration. **A.** Treatment regimen. **B-D** Immunofluorescent imaging (**B**) and quantification (**C, D**) of collagen 1 fibers and the basement membrane (BM) components laminin and perlecan in WT, KO and KO+PBS mouse cornea. The graphs in **C** and **D** show expression levels of laminin and perlecan in the central corneal BM. **E** and **F.** Distribution of collagen 1 fiber thickness (**E**) and alignment angle in relation to the BM (**F**) in the corneal stroma of each group. **G.** Central corneal stromal thickness in each of the 3 groups. **H.** Heat map highlighting representative genes associated with extracellular matrix (ECM) in the cornea that were altered 24 hours post treatment. NUC = nuclei. Scale bar = 50 μm *p < 0.05; ***p < 0.001; Data in **B, C** and **F** were subjected to a one-way analysis of variance with a post-hoc Tukey’s test. Data in **D** and **E** were subjected to a Kolmogotrov-Smirnov test. Each dot in the bar graph represents a biological replicate. Error bars represent standard deviation. n > 4 mice per group.

We further defined alterations in gene signatures associated with the stroma and BM in the 3 conditions by performing bulk RNA sequencing (RNAseq) of corneas isolated 24 hours (hr) after lubrication. Major transcriptional differences were identified between each condition, as shown by principal component analysis (PCA), consistent with extensive changes in tissue structure and function (Supplementary Figure S1C). In support of stromal regeneration, KO corneas treated with PBS exhibited a significant upregulation of gene transcripts essential to stromal ECM and basal epithelial BM integrity as compared to untreated KO (Figure 1H and Supplementary Figure 1D). These included the BM-associated genes collagen 4 (*Col4a3* FC = 194, p=1E-28; *Col4a4* FC = 74.5, p =1E-21), laminin (*Lama2* FC = 3.5, p <0.05), nidogens (*Nid1* FC = 2.7, p = 1E-07, and *Nid2* FC = 6.6, p < 0.01) and the stromal ECM-related genes keratocan (*Kera* FC = 4.6, p < 0.01), lumican (*Lum* FC = 7.4, p =1E-5) and fibronectin (*Fn1* FC = 2.3, p <0.01). In addition, lubrication resulted in the downregulation of gene transcripts for proteins involved in ECM degradation e.g., metalloproteinase (*Mmp)9* (FC = 7.36, p < 0.01), a clinical indicator for dry eye disease, while others involved in blocking degradation were upregulated, e.g., metalloproteinase inhibitor 3 (*Timp3*, FC = 26.9, p =1E-11). Lubrication also elevated gene expression of several factors known to actively contribute to wound healing and tissue repair including fibroblast growth factor 1 (*Fgf1*), insulin growth factor 2 (*Igf2*), and thrombospondin-1 (*Thbs1*) and increased expression of a set of crystallin genes (e.g. *Cryaa, Crygb, Crygd*) that contribute to corneal transparency and are lost in dry eye disease (15) (Figure 1H).

Thus, these data suggest PBS is sufficient to promote the restoration of stromal and BM integrity as well as cell-matrix interactions critical to the maintenance of corneal transparency and visual acuity.

### Lubrication reveals keratocytes to possess extensive plasticity and to undergo regenerative transformation with treatment

Single cell RNA sequencing (scRNAseq) has identified heterogeneous stromal keratocytes in healthy human and murine corneas under resting conditions(7, 16). However, despite keratocytes being known to shift from a resting to an activated state following injury, the changes in their cellular profiles under pathological or healing conditions, and the mechanisms driving these changes, remain unknown. To investigate keratocyte heterogeneity in the 3 conditions, the dynamic transcriptomic signatures of these cells during their transition, and the associated changes in signaling pathways, we analyzed tissue via single-nuclear RNA sequencing (snRNAseq). Nuclei were extracted from whole corneas (n=5, 10 corneas per group) and subjected to snRNAseq using the 10x genomics platform (Figure 2A). We identified 3153 WT, 5123 KO and 3558 KO+PBS keratocytes enriched in ECM transcripts including collagen 1(*Col1*), decorin (*Dcn*), and lumican (*Lum*). Cell filtering and unbiased clustering (R package Seurat) defined 10 keratocyte populations (Figure 2B-C and Supplementary Figure S2A), with each treatment group being composed of multiple distinct cell clusters. WT cornea was composed of 3 clusters (cluster 2, 4, 8), with clusters 4 (28%) and 8 (10%) being enriched in ECM component and remodeling genes *Col6a1* and *Col1a1* and metalloproteinase *Mmp3* and *Col3a1*, respectively, suggestive of actively remodeling keratocytes. In comparison, cluster 2 (62%) was marked by low levels of *Col1a1*, the major collagen within the bulk of the fibrils, suggesting these cells are “resting” keratocytes that have a key role in regulating stromal architecture.

**Figure 2.**
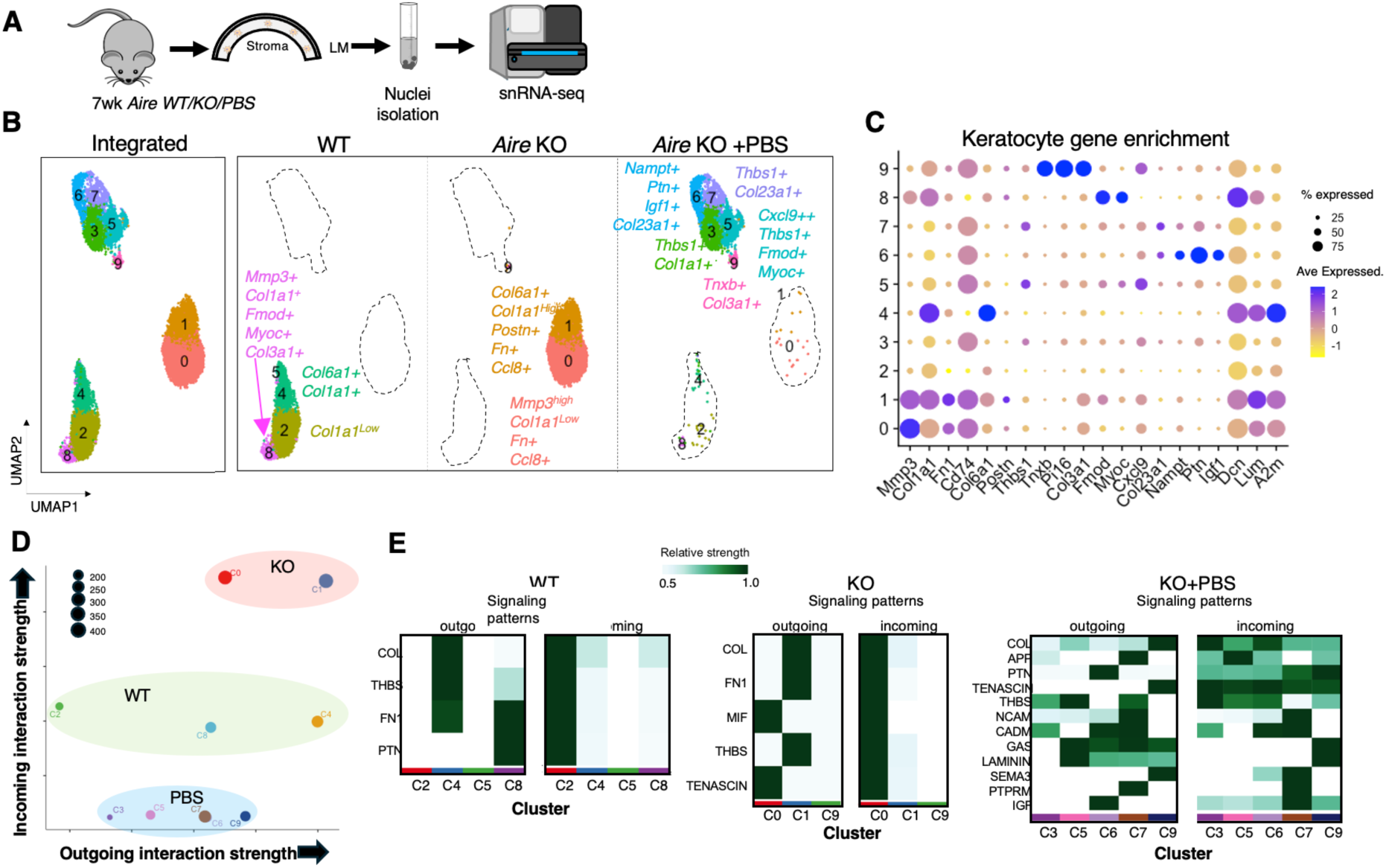
Keratocytes in the diseased corneal stroma exhibit striking plasticity in response to disease and lubrication. **A.** Workflow for cornea collection, nuclei isolation, and sequencing. **B.** The Integrated Stroma UMAP identifies 11,835 keratocytes from all 3 samples distributed into 10 clusters by unsupervised clustering. Individual UMAPs showing the distribution of keratocytes within each condition and specific marker genes for each cluster. **C.** Dot plot of the top marker genes for each identified cluster. The size of the dot corresponds to the percentage of cells within the cluster expressing the gene. The average expression level for the listed genes is indicated by the color. **D**. Scatter plot visualizing dominant senders and receivers of cellular signals in a 2D space. x-axis and y-axis are respectively the total outgoing and incoming communication probabilities associated with each cluster for each group. **E**. Heatmaps identify signals contributing the most to outgoing or incoming signaling of the specific keratocyte clusters for each condition.

In comparison, the KO keratocytes existed as two clusters (cluster 0, 1) that were dramatically different to the WT. Both clusters were enriched in inflammatory and injury associated genes (e.g., *Ccl8, Cd74*, *and Slpi,* Figure 2B-C) as well as in fibronectin (Fn), a factor known to be highly upregulated with chronic injury (17). Cluster 0 (61%) was highly enriched in *Mmp3*, a proteolytic protein overexpressed with chronic damage to promote matrix degradation and inflammation (18). Cluster 1 (39%) was highly enriched in periostin (*Postn),* a marker of pro-fibrotic fibroblasts in chronic injury conditions (19)*, and Col1a1, Col6a1,* proteins that are well known to contribute to fibrosis when overexpressed during aberrant wound healing (19–21). These data suggest chronic injury provokes keratocytes to transition into two destructive, fibrotic cell populations with each cluster likely possessing a unique role in stromal disruption and impaired cornea regeneration.

Strikingly, topical lubrication further modified the keratocyte cell states resulting in five unique clusters (clusters 3, 5, 6, 7 9) that were specifically enriched in genes associated with wound repair. Clusters 3 (36%), 5 (25%), and 7 (15%) were enriched in *Thbs1, a* matricellular protein released and/or secreted following tissue injury that is typically associated with ECM organization (22) and inhibiting angiogenesis (23). Clusters 6 (18%) and 7 were both enriched in *Col23a1*, a non-fibrillar collagen involved in cell-cell and cell-matrix adhesion (24, 25). Cluster 6 was also enriched for *Igf1*, which has been shown to support corneal epithelial wound healing (26), and pleiotrophin (*Ptn),* a factor with neurotrophic activity that promotes nerve regeneration and is actively engaged in bone marrow stromal cell-mediated regeneration (27). Cluster 9 (4%) was strongly enriched in stem cell-associated factors, including the matrix proteins *Tnxb* and *Col3a1*, which provide structural support and regulate tissue flexibility and elasticity (28,29). These proteins have been shown to be expressed by limbal keratocytes (30, 31). Additionally, the peptidase inhibitor *Pi16,* which is highly expressed in stromal/fibroblast stem-like cells across various tissues (32) was also enriched, suggesting that Pi16 may also mark limbal keratocyte stem cells. Consistent with lubrication providing therapeutic benefit, relative to the untreated KO there were no cell clusters enriched in pro-fibrotic or chronic injury markers such as *Col6a1* or *Mmp3* in the lubricated cornea, supporting the pro-regenerative state of the tissue (Figure 2B-C).

Tissue homeostasis and repair are complex processes involving activation of various signaling pathways and intercellular coordination. Through analysis of intercellular communication networks via CellChat (33), we identified differences in cell signaling between clusters in each condition that were consistent with the marker genes. In the WT cornea, clusters 4 and 8 showed active signaling to the surrounding cells such as COLLAGEN, THBS and FN1 signaling, while cluster 2 was the main receiver of cellular signaling (Figure 2B,D-E). The two KO cell clusters exhibited the greatest levels of active signaling networks in both incoming and outgoing signaling among all the keratocyte clusters including inflammatory signaling-MIF and TNX that were not seen in the WT (Figure 2D). Comparison of the intercellular communication networks between the two KO clusters indicated cluster 0 is the main receiver of incoming cellular signaling as well as the source of outgoing inflammatory signaling-MIF, suggestive of an activation of an autocrine inflammatory loop within these keratocytes (Figure 2E). CellChat analysis further revealed that the 5 unique clusters as key sources of THBS, PTN, IGF, and TENASCIN signaling in the lubricated keratocytes (Figure 2E), supportive of their reparative/regenerative role. The observed increase in complexity of the intercellular communication network further illustrates reparative actions of the keratocytes in the lubricated cornea (Figure 2E). Together, our data reveal the dynamics of keratocyte cell state under homeostatic, diseased and lubricated settings and that lubrication is sufficient to produce a reparative keratocyte phenotype.

We next validated the different clusters of keratocytes and identified their spatial locations within the cornea through in-situ hybridization (RNAscope HiPlex) and immunofluorescent analysis (Figure 3). In the WT, we found that the abundant *Col1a1* enriched cells (cluster 2) were distributed throughout the corneal stroma, consistent with the corneal-wide array of collagen fibers and their known enrichment in COL1 (Figure 3A-B). Clusters potentially involved in ECM remodeling (cluster 4, 8), were located within the limbal and peripheral regions of the cornea. Specifically, *Col6a1+* cells (cluster 4) were most abundant in the peripheral cornea (Figure3A-B and Supplementary Figure S3A), suggesting a potential role of these cells in forming distinct networks of microfibrils (34). A small proportion of WT keratocytes (Cluster 8) enriched in fibromodulin (*Fmod),* a proteoglycan critical to scarless wound healing in skin, and *Mmp3*, a key player in ECM remodeling, were concentrated at the limbal region (Figure 3B), suggesting this population to be involved in maintaining stromal architecture during homeostasis.

**Figure 3.**
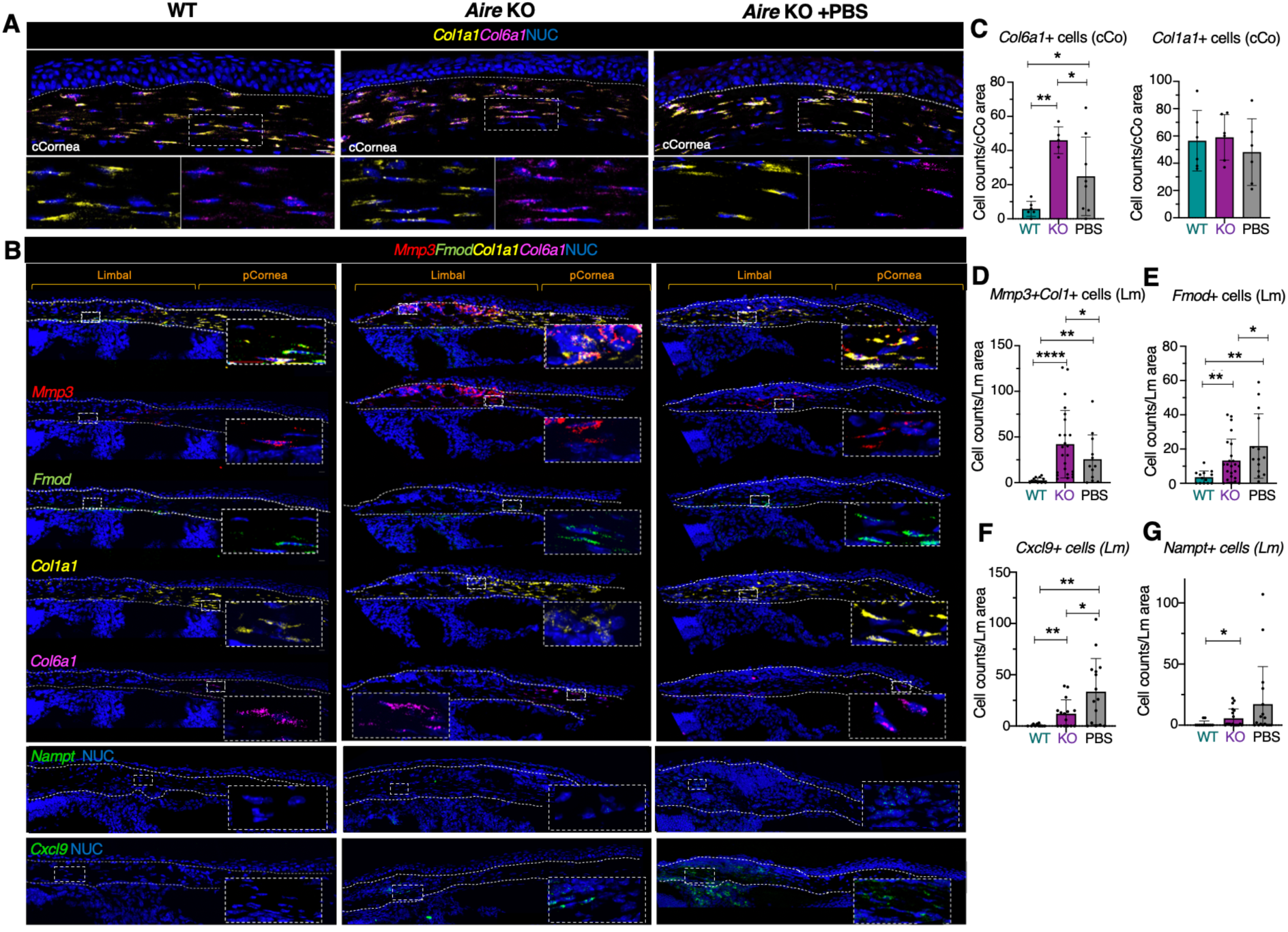
Lubrication promotes a reparative response in the diseased stroma by reducing cells associated with excessive ECM remodeling and enriching for those promoting wound healing. **A-B.** In-situ hybridization analyses of marker genes revealing spatial location of the keratocyte clusters in the central cornea (**A**, cCornea) and peripheral cornea (pCornea) and limbal region (**B**). Boxes show high magnification of stromal cells within the stromal region. Dotted lines outline the upper and lower borders of the stromal compartment. **C-G.** Quantification of the number of keratocytes expressing high transcripts levels of target genes in the 3 groups. *p < 0.05; **p < 0.01; ***p < 0.001; **** p < 0.0001; Data in C-G were subjected to a one-way analysis of variance with a post-hoc Tukey’s test. Each dot in the bar graph represents a biological replicate. Error bars represent standard deviation. n > 4 mice per group. Scale bars = 50 μm.

Similar to WT, cells enriched in *Col1a1* and *Col6a1* in the KO were distributed across the peripheral and central cornea with very few *Col6a1+* cells located in the limbal region (Figure 3A-B). However, the number of *Col6a1*^high^ keratocytes in the KO was significantly greater than WT while numbers of *Col1a1*^high^ keratocytes remained relatively unchanged (Figure 3A, C). Although COL6 is a nonfibrillar collagen that plays an important role in wound healing and tissue repair (35), however, excessive production of COL6 is often associated with tissue fibrosis (20) suggesting that an expansion of this population in the KO impairs effective wound healing. Similar to the WT keratocytes enriched in *Mmp3* and *Fmod* (cluster 8), KO *Mmp3+* (cluster 0) and *Fmod+* keratocytes (cluster 1) were specifically found in the limbal region in close association with infiltrating immune cells (Figure 3B). Strikingly, the number of these cells at this location dramatically increased with disease (Figure 3B, D-E), consistent with the size of *Mmp3*-enriched cluster 0 and *Mmp3* expression level (Figure 2B-C), suggesting excessive ECM remodeling and aberrant wound healing were taking place in the limbal region.

We next validated the presence of the potentially regenerative clusters enriched in the lubricated cornea. *Thbs1+* keratocytes (clusters 3, 5 and 7), were located throughout the corneal stroma, consistent with THBS1 being essential for avascularity and transparency as well as accelerated wound healing in the corneal epithelium(23, 36) (Supplementary Figure S3B). Cells enriched in nicotinamide phosphoribosyltransferase (*Nampt*), a protein shown to protect cells from apoptosis (37, 38) (cluster 6), and chemokine (C-X-C motif) ligand 9 (*Cxcl*9), an immune modulator (39) ( cluster 5), were localized to the limbal stroma (Figure 2B,C and Figure 3B, F, G). In addition, in the lubricated cornea, the population of *Col6a1*^high^ and *Mmp3*^high^ keratocytes was greatly reduced (Figure 3A-D) while *Fmod*^high^ expressing keratinocytes were increased (Figure 3B, E), potentially indicating the reversal of excessive ECM remodeling and the transition of these keratocyte populations to a reparative state. In summary, our data reveal the plasticity of keratocytes and their extensive remodeling in response to disease and treatment with lubrication. Moreover, we demonstrate a regenerative function of topical lubrication in promoting keratocyte transition from a disease-associated, destructive state to a reparative cell state.

### Lubrication promotes optimal collagen production while dampening keratocyte inflammation

To gain a deeper understanding of the heterogeneity and intercellular relationships within each condition, we examined the WT, *Aire* KO, and *Aire* KO + PBS datasets separately and further sub-clustered the keratocytes. We identified four unique WT clusters, each representing a keratocyte population with specific functions (Figure 4A-C and Supplementary Figure S4A). For instance, cluster 0, the largest cluster constituting more than 50% of the keratocyte population, represented the resting keratocytes with low level of decorin and collagen transcripts; cluster 1 (23.57%) was highly enriched in neuronal genes including *Erbb4, Slit3, Cped1, and Auts2*, suggesting their interaction with nerves; cluster 2 (16.64%) was enriched in ECM genes, indicating their potential contribution to ECM remodeling during corneal homeostasis; cluster 3 (9.56%) was enriched in the limbal markers, *Fmod* and *Mmp3*, indicating they are exclusively located in the limbal region (Figure 4B-C and Supplementary Figure S4A). This cluster was also enriched for *Igfbp5*, a factor known to promote deposition of collagen and fibronectin by fibroblasts (40, 41). Potential stem cell marker *Pi16,* as well as *Cd34,* a factor expressed by putative keratocyte progenitors in the human cornea (42) and expressed in the limbal region of the murine corneal stroma (Supplementary Figure S4B), were enriched specifically in a small proportion of cluster 3, supporting the existence of a limbal keratocyte stem cell (LSC) population within this cluster (Figure 4B).

**Figure 4.**
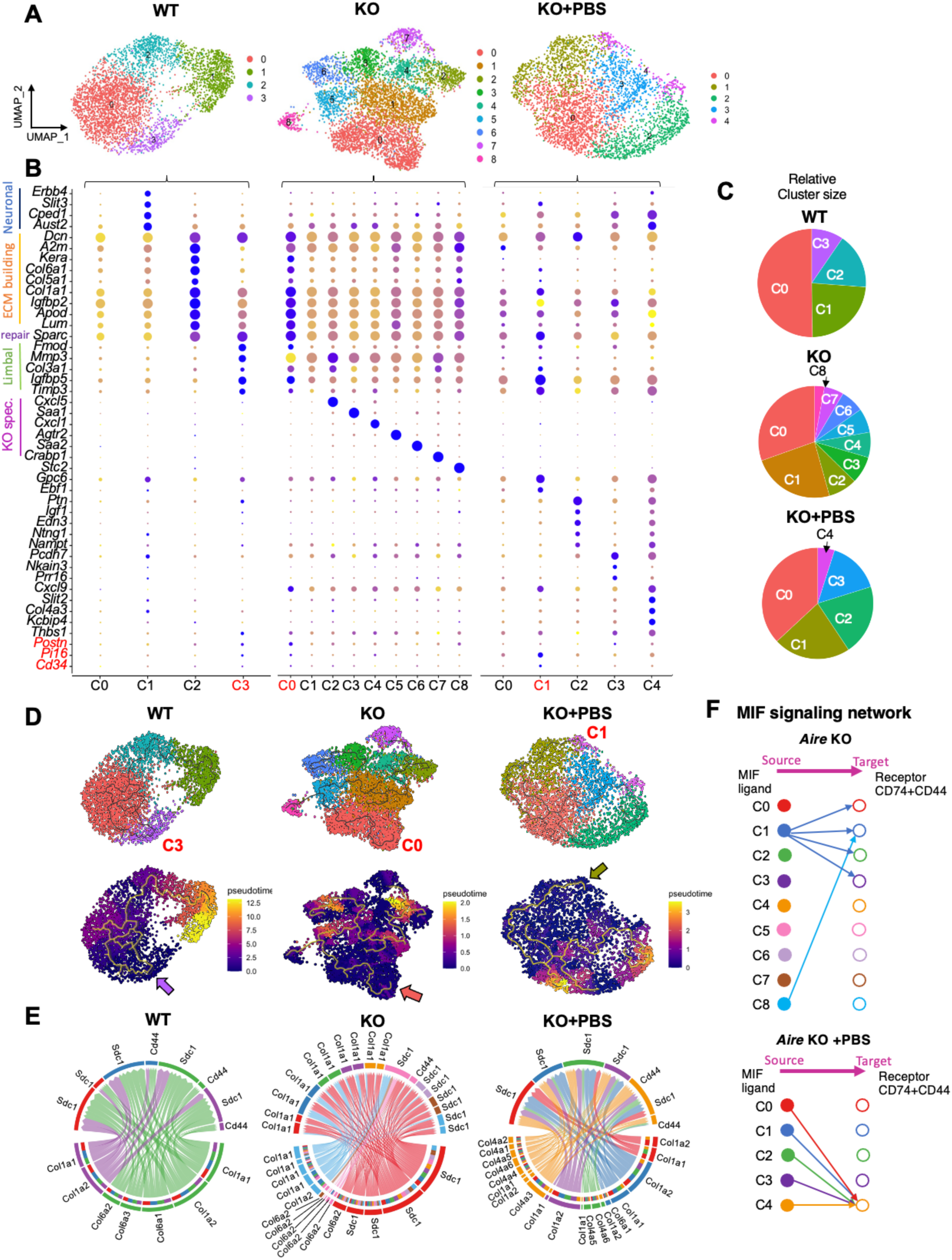
Leveraging the keratocyte plasticity to enhance repair by promoting optimal collagen production and dampening keratocyte inflammation. **A,B.** UMAP (**A**) and respective dot plot (**B**) highlighting the heterogeneity of keratocytes within each condition and the top marker genes for each identified cluster. The size of a dot corresponds to the percentage of cells within the cluster expressing the gene. The average expression level for the listed genes is indicated by the color. **C.** Pie chart summarizing the proportion of keratocytes in each cluster within the different groups. **D.** Monocle trajectory inference and pseudotime analysis showing keratocyte transitional states within each condition. Dark purple represents an earlier stage while yellow represents a later time point. The arrow points to the initiating cell location. **E.** Chord plots of collagen signaling pathway network showing ligand and receptor interactions between keratocyte clusters within different conditions. **F.** Comparison of MIF signaling pathway network and ligand/receptor interactions between keratocyte clusters in *Aire* KO vs. PBS.

In contrast to the WT, we identified nine different clusters within the inflamed and pro-fibrotic stromal keratocytes of the *Aire* KO cornea (Figure 4A) The resting keratocyte, cluster 1, was dramatically reduced to 23.85% compared to WT (50.23%) while the ECM remodeling cluster 0 was significantly increased to 30.45% compared to 9.56% in WT (Figure 4A, C and Supplementary Figure S4A). Cluster 0 was also enriched in *Sparc*, a gene associated with corneal wound healing and regeneration (43). Interestingly, the limbal keratocyte markers were enriched in both clusters 0 (i.e., *Fmod, Postn, Igfbp5*) and 2 (*i.e., Mmp3*) while the LSC markers, *Pi16* and *Cd34*, were predominantly limited to cluster 0. Notably, neuronal-associated genes enriched in the WT cornea were absent in the KO keratocytes while clusters 2 through 8 were each uniquely enriched in a specific marker gene that defined their immune modulatory function (Figure 4B, KO dot plot). For instance, *Cxcl5* (cluster 2) plays a role in recruiting lymphocytes and eosinophils during an immune response (44); *Saa1*(cluster 3), induces inflammatory gene expression in fibroblasts (45); *Cxcl1* (cluster 4) is a chemoattractant cytokine that primarily recruits neutrophils (46); *Agtr2* (cluster 5) mediates tissue fibrosis signaling in activated fibroblasts (47); *Saa2* (cluster 6), amplifies innate immune responses (48); *Crabp1* (cluster 7), marks wound-healing fibroblast (49); and Stc2, which was exclusively enriched in the smallest population (cluster 8 (2.86%)), is known to inhibit the recruitment and regeneration of CD8+ T cells (50), indicative of an anti-inflammatory role.

Intriguingly, lubrication reduced keratocyte heterogeneity to five distinct clusters, with an increase in the resting keratocyte cluster (C0, 37%) compared to that of the untreated KO cornea (C0, 24%, Figure 4A-C and Figure S4A). Additionally, the enrichment of neuronal associated markers noted in WT mice was restored with PBS, suggesting the re-establishment of nerve-keratocyte interactions. Cluster 1 enriched in signature genes for ECM remodeling, including Kera, Lum, and Col6a1, was reduced compared to the untreated *Aire* KO (PBS: 23.41% vs. KO: 30.45%) while clusters 2 through 4 (totaling 40%) were enriched in marker genes associated with wound healing and regeneration, including *Ptn, Igf1, Thbs1*, and *Col4a3*. *Col4a3* is one of the main components of the corneal basement membrane expressed predominantly by keratocytes, thereby supporting our finding that lubrication alone enhances BM regeneration in the *Aire* KO cornea (Figure 1B). Finally, limbal keratocyte markers *Mmp33, Fmod,* and *Postn* were enriched in C1 and C3, whereas the limbal stem cell markers *Pi16* and *Cd34* were restricted to the ECM remodeling cluster (C1), along with the wound healing gene, *Sparc*. Together, these data further highlight dry eye induction of specific inflammatory cell types and the active transition of these keratocytes to a reparative state with simple lubrication.

To investigate the relationship among keratocyte subclusters under homeostatic, disease, and regenerative conditions, we conducted lineage trajectory analysis using monocle 3 (Figure 4D). Predicted trajectory graphs revealed the remarkable plasticity of stromal keratocytes, showing their ability to transition between different functional cell states. For instance, resting keratocytes in *Aire* KO C1 shift into various cell states, such as the ECM-remodeling cluster 0 (ECM builder) via cluster 5, or the ECM-degrading cluster 2 via cluster 4 (Figure 4D, upper). Pseudotime analysis of the keratocytes, with the starting cluster being the stromal limbal stem cells (arrows) (Figure 4D, lower) revealed a linear trajectory ending with the neuronal-associated cluster in the WT cornea (C1), placing it at the later differentiated stage relative to the other clusters. However, no clear trajectory path from early to late differentiation was found in the KO or PBS conditions, further supporting the notion of keratocyte plasticity during wound healing.

To delineate and compare intracellular communication among the clusters under each condition, we conducted Cellchat analysis with the goal of identifying the source and target of signaling pathways within the keratocytes via ligand and receptor interaction. Collagen signaling was the most enriched pathway for keratocytes in all three conditions, yet the type of collagen produced by the keratocytes and its level of expression was different (Figure 4E). In the WT and untreated *Aire* KO, all keratocytes expressed collagen receptors *Cd44* and *Sdc1*, while the ECM building cluster (C2 in WT and C0 in KO) was the main producer of collagens 1 and 6. In the untreated KO, two other clusters (C5, C8) were also making collagens 1 and 6 contributing to an overall increase in collagen signaling. This increase in *Col6a^high^* keratocytes was validated in the *Aire* KO cornea (Figure 3A,C), supporting the pro-fibrotic environment. In stark contrast, all keratocyte clusters in PBS-treated cornea were producing collagens, with an especially notable downregulation of *Col6* and upregulation of *Col4,* which is essential for basement membrane rebuilding, thereby illustrating the transition of keratocytes from a destructive to reparative state. All PBS keratocytes expressed collagen receptor *Sdc1* with only one cluster (C4) expressing both *Sdc1* and *Cd44*. CD44 is a multi-functional cell surface adhesion receptor involved in a range of cellular processes including cell adhesion, migration, cell signaling, inflammation, and wound healing (51). Its ability to bind multiple ligands makes it crucial in regulating cell behavior in response to environmental changes, such as those that occur during an immune response. As such, the reduction of *Cd44* expression with lubrication may signify reduced keratocyte inflammation (Figure 4E).

One of the most intriguing findings was the profound enrichment of the Macrophage migration Inhibitory Factor (MIF) pathway in *Aire* KO keratocytes that was nearly absent in response to PBS treatment (Figure 4F). MIF is known to be upregulated in inflamed tissues and signals through two essential co-receptors, CD44 and CD74, both of which are required for signaling (52,53). In the untreated *Aire* KO, the MIF1 ligand was produced by C1 (∼24%) and C8 (∼3%), while C0-C3, which comprise 60% of the cells, served as targets, expressing both MIF1 co-receptors CD74 and CD44. In comparison, while all PBS-treated keratocytes expressed the MIF ligand, only the small cluster C4 (4%) expressed both co-receptors (Figure 4F), indicating downregulation of MIF signaling and supporting the idea that lubrication dampens keratocyte inflammation. Notably, PBS-treated keratocytes in C4 were also enriched with neuronal- and BM- associated genes, suggesting the cells were reparative. Indeed, in addition to its role as an inflammatory mediator, MIF possesses wound healing functions through regulating many novel repair/inflammation-associated gene targets e.g., *Col1a1, Cxcl10*, and *Thbs1*, and as such, may contribute to the regenerative properties of this cell cluster. Overall, our results indicate keratocytes exhibit significant plasticity in adapting to environmental cues.

### Keratocytes display distinct transcription factor (TF) profiles that correlate with inflammatory and reparative processes

Next, we explored potential TFs involved in regulating keratocyte cell state transition in disease and during tissue repair using single-cell regulatory network inference and clustering (SCENIC), a computational method that predicts critical regulators and their direct target genes (54). SCENIC identified 114 TFs to be highly enriched in keratocytes and created an unbiased clustering of cells based on their specific TF profiles. From cell clusters identified by gene expression analysis (Seurat, Figure 5A, left), 3 major TF-defined cell clusters emerged, with each TF cluster being specifically composed of keratocytes (single dots) from either the WT (clusters 2, 4, 8), KO (clusters 0 and 1) or KO+PBS (clusters 3, 5, 6, 7, 9), thus indicating high similarity of TF profiles within each treatment group (Figure 5A, right).

**Figure 5.**
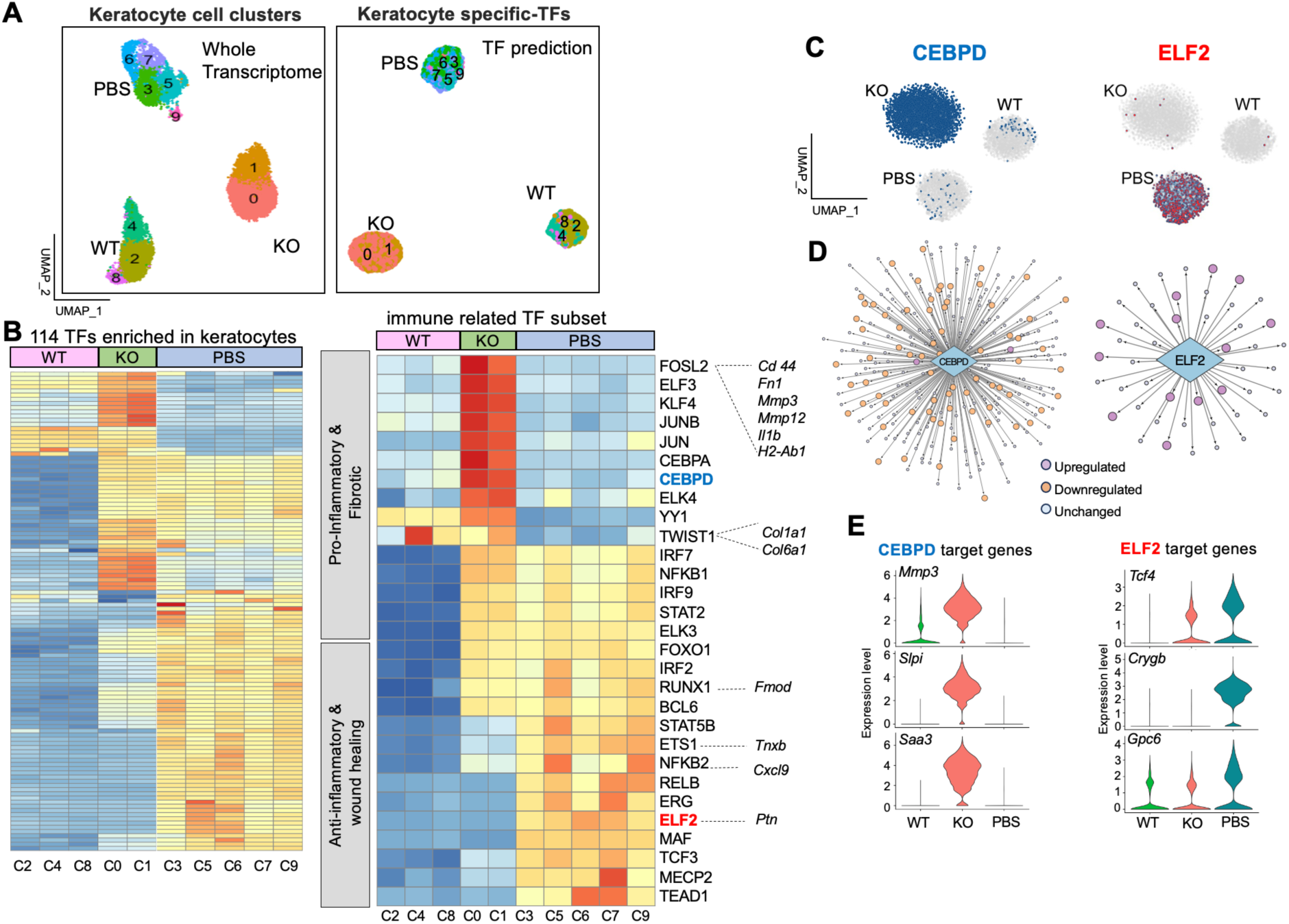
Keratocytes display distinct transcription factor (TF) profiles that correlate with alteration in inflammatory and reparative processes. **A.** UMAP plot showing keratocytes from WT, KO and KO+PBS distributed into 3 major clusters. Left UMAP shows clustering based on the whole transcriptome and the right shows unsupervised clustering based on SCENIC TF prediction. **B.** Heatmaps of AUC scores of enriched TFs in all keratocyte clusters, as predicted by SCENIC. Left heatmap is all 114 predicted TFs enriched in the keratocytes. Right highlights a subset of predicted inflammation and wound healing associated TFs along with examples of target genes. Color coded from blue to red indicates increased regulon activity. **C.** t-SNE plots showing the TFs CEPBD and ELF2 enriched in KO and KO+PBS keratocyte clusters, respectively. **D**. The extent of gene regulatory networks regulated by CEBPD and ELF2. Each line represents a target pathway, with circles indicating pathways that are up- or down-regulated or unchanged. **E.** Violin plots of representative target genes for CEBPD and ELF2 comparing all 3 groups.

The completed heatmap revealed all regulons from the 114 TFs identified to be greatly enriched in the keratocytes. We found high similarity amongst the clusters belonging to the same treatment group, however, each individual cluster had its own unique TF profile (Figure 5B, left). As would be predicted, compared to WT and PBS samples, the *Aire* KO clusters were enriched in inflammation-associated TFs (eg. FOSL2, JUN, KLF4, CEBPD, and TWIST1) (Figure 5B, right) that regulate fibrosis, angiogenesis and/or fibroblast inflammatory processes (55–57). This included FOS-Like 2 (FOSL2), which regulate a cassette of pro-inflammatory target genes, such as *Il1b*, *Mmp*3, *Fn1* and *H2-Ab1,* and Twist Family BHLH Transcription Factor 1 (TWIST1), which regulates ECM remodeling genes including *Col1a1* and *Col6a1*(SCENIC predictions, Figure 5B, right). Lubrication resulted in reduced expression of multiple target genes within inflammatory TF regulons. For example, of the 234 target genes of CEBPD enriched in *Aire* KO keratocytes, as shown in the t-SNE plot (Figure 5C), 72 were downregulated with PBS treatment (Figure 5D), including ECM remodeling genes *Mmp3, Mmp12, Lum, Col1a2,* and *Col3a1*, as well as fibroblast inflammatory genes *Slpi* and *Saa3* (Figure 5D-E), supporting the notion that prolonged dryness drives chronic keratocyte activation. Furthermore, analysis of the KO+PBS dataset revealed the enrichment of TFs strongly associated with anti-inflammatory and wound healing responses (eg. NFKB2, ETS1, MECP2, ELF2, MAF). Intriguingly, target genes were found to mark specific cell clusters enriched in the lubricated cornea (Figure 5B). For example, *Cxcl9*, *Tnxb*, and *Ptn* which are regulated by the TFs NFKB2 and the ETS family transcription factors ETS and ELF2 marked cell clusters 5, 9 and 6, respectively. ELF2 is an inhibitor of myofibroblast activation (58) and regulates a total of 47 target genes, 13 of which are upregulated with PBS. These included genes associated with corneal transparency (*Crygb)* (59), hedgehog mediated-corneal wound healing (*Gpc6)* (60), and tissue regeneration (fibroblast marker *Tcf4)* (61) (Figure 5C-E). Finally, we found a number of inflammatory TF regulons to be unchanged with/out lubrication such as IRF7, NFKB1 and STAT2, indicative of a persistent pro-inflammatory environment, suggesting that lubrication facilitates stromal repair despite keratocyte inflammation.

Together, these data suggest each condition utilizes a distinct set of TFs, pro-inflammatory/pro-fibrotic and anti-inflammatory/pro-repair, to regulate keratocyte activation and the transition from destructive to reparative cell states.

### Lubrication rescues stromal architecture via inhibition of an IL1 autocrine loop

To define the mechanism by which lubrication rescues stromal architecture, we performed gene ontology (GO) analysis of our global and single nuclei RNAseq comparing WT, KO and KO+PBS. Comparison of *Aire* KO vs WT revealed the *Aire* KO keratocytes to have a significant enrichment of pathways associated with interleukin signaling, and more specifically IL1β (Figure 6A and Supplementary Figure S5A). IL1β has a demonstrated central role in a number of autoinflammatory diseases and is a mediator of many local and systemic features of inflammation including activation of MIF pathway by upregulating MIF ligand expression (62). It also triggers a positive, inflammatory feedback loop that amplifies the IL-1 response through autocrine or paracrine signaling (63, 64). We previously showed that *Il1b* and its primary receptor *Il1r1* are upregulated in *Aire* KO cornea, with *Il1b* being highly expressed by the epithelium and *Il1r1* by both epithelial cells and keratocytes (65). snRNA sequencing showed an increase in both *Il1r1*^high^ and *Il1b-expressing* keratocytes in the *Aire* KO cornea compared to the wild type (Figure 6B). This outcome is supported by our immunofluorescent analysis revealing a significantly higher number of IL1β^high^ IL1R1 expressing keratocytes in the *Aire* KO stroma than WT and lubricated KO corneas (Figure 6C-D). Although the number of *Il1r1*^high^ keratocytes was not reduced with lubrication compared to untreated KO (WT= 0%, KO=4%, PBS=5%), lubrication led to a decrease in the number of *Il1b*-expressing keratocytes (WT=- 0.3%, KO=10.7%, PBS=2.7%) (Figure 6B-D). Our global RNAseq analysis also shows a significant reduction in *Il1b* in the KO cornea treated with PBS (PBS vs KO. FC=6.2, p<0.01) (Figure 1H), suggesting lubrication inhibits IL1β signaling through downregulating *Il1b* production in keratocytes.

**Figure 6.**
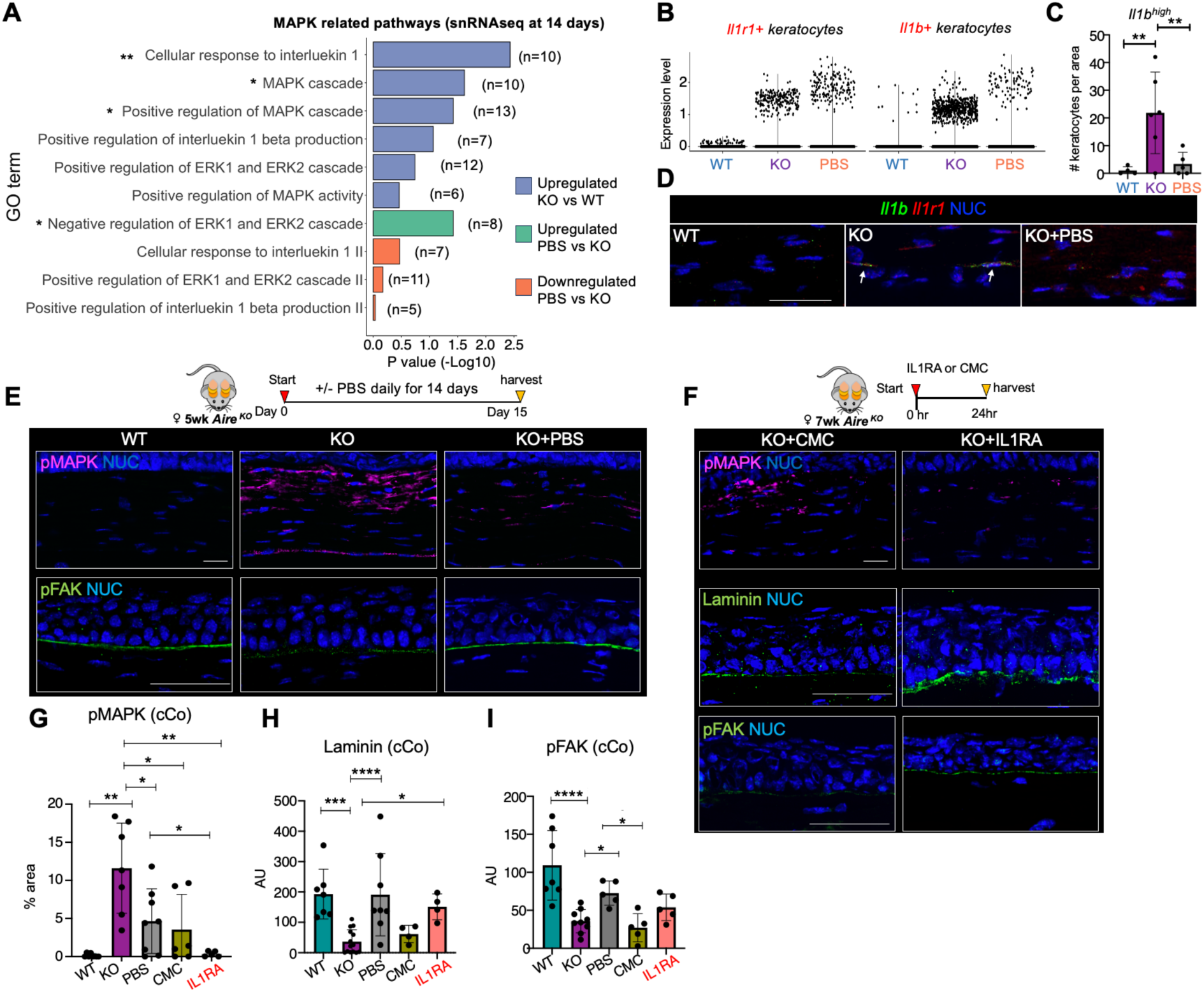
Lubrication rescues stromal architecture via inhibition of IL1B-IL1R1-MAPK signaling. **A.** Gene Ontology (GO) analysis of snRNAseq datasets. n= number of significant differentially expressed gene in the pathway. **B.** Violin plots comparing *Il1r1* and *Il1b* expressing cells in each WT, KO, and KO+PBS groups. Each dot represents a cell. **C-D**. Immunofluorescent analysis of cells expressing IL1B, (**C**) and quantification of *Il1b ^high^*keratocytes (**D**) in the cornea stroma of the 3 groups. **E.** Schematic of topical treatment regimen for 14 days starting at 5 wks of age. Immunofluorescent analysis of phosphorylated (p) MAPK (upper panel) and pFAK (lower panel) in WT, KO and KO+PBS stroma. **F.** Schematic of topical treatment regimen in 7wk old mice. Immunofluorescent analysis of pMAPK, laminin and pFAK in WT, KO and KO+PBS stroma at 24 hr. **G-I.** Quantification of expression levels of pMAPK (**G**), laminin (**H**) and pFAK (**I**) in the central cornea (cCo). NUC = nuclei. Scale bar = 50 μm *p < 0.05; **p < 0.01; ***p < 0.001; **** p < 0.0001; Data in C, F, G and H were subjected to a one-way analysis of variance with a post-hoc Tukey’s test. Each dot in the bar graph represents a biological replicate. Error bars represent standard deviation. n > 4 mice per group.

Signal propagation by IL1β mainly depends on mitogen-activated protein kinases (MAPKs specifically ERK), a signaling pathway that we found to be significantly upregulated in keratocytes with disease (Figure 6A). Surprisingly, our single cell analysis showed lubrication alone was sufficient to significantly downregulate this signaling cascade at 14 days of treatment (Figure 6A). This reduction of the IL1β and MAPK signaling at the single cell level was further supported by global RNAseq analysis after 24 hours of lubrication (Supplementary Figure S5A), suggesting IL1β-MAPK signaling contributes specifically to keratocyte transition to an inflammatory cell state in the desiccated cornea and that this transition can be suppressed by lubrication. Coincidentally, pMAK is also a downstream target of MIF signaling (66). Therefore, a reduction in global IL1b levels could lead to the downregulation of the MIF signaling pathway (Figure 4F) and its downstream target, pMAK. This suggests that IL1b-MAPK signaling in KO keratocytes may be either MIF-dependent or independent. Our data further indicate that lubrication reduces IL1-mediated inflammation through the downregulation of *Il1b* production by keratocytes.

To validate these outcomes we performed immunofluorescent analysis of MAPK activity in *Aire* KO, WT, and KO+PBS cornea. pMAK expression was extensively elevated in the corneal stroma of the *Aire* KO compared to the WT, while it was dramatically reduced by topical lubrication of the cornea for 14 days (Figure 6E, G), thereby confirming our transcriptomic findings. To determine if inhibition of IL1β signaling was sufficient to rescue the stromal architecture, we blocked the receptor for IL1β, interleukin-1 receptor 1 (IL1R1). *Aire* KO corneas treated with an IL1R1 antagonist (IL1RA) for 24hrs showed a robust reduction in pMAK compared to untreated Aire KO or vehicle control (Carboxymethyl Cellulose (CMC)), with levels resembling the WT (Figure 6E-G). As expected for a lubricant, CMC showed some reduction in pMAPK at 24 h (Figure 6F-G), complementing our global RNAseq analysis showing negative regulation of ERK1 and ERK2 (MAPK) cascade at 24hr of PBS treatment (Supplementary Figure S5A).

Next, we assessed the response of the desiccated cornea to blocking IL1R1 signaling for 24 hrs on BM and stromal integrity. The BM and stroma were evaluated by immunostaining for laminin deposition, active focal adhesion kinase (FAK), a mechanosensor phosphorylated (p) in response to ECM stiffness, and collagen fiber alignment. Despite the short time frame, 24 hrs of treatment with the IL1R1 antagonist, but not with CMC alone, resulted in a dramatic increase in laminin deposition at the basal epithelium (Figure 6F, H). Although CMC alone (a common ingredient found in over-the-counter ocular lubricants used by dry eye patients) did not restore the BM, global RNAseq analysis following 24h of lubrication revealed an upregulation of BM associated genes such as *Lama2*, *Col4a3*, and *Nid1,* indicating the activation of BM regeneration occurs within this short time frame (Figure 1H). However, longer periods of lubrication, such as 14 days of PBS treatment, did restore the BM. Similarly, pFAK in the central cornea was restored to WT levels after 14 days of lubrication (Figure 6E, I). There was also an increase in pFAK levels, albeit not significant, within 24hrs of IL1RA treatments (Figure 6F, I) suggestive of an improvement in BM assembly that correlated with laminin deposition. Given that laminin is synthesized by both basal epithelial cells and keratocytes, and pFAK accumulates at the adhesion points along the BM, restoring BM implies re-establishing the epithelial-matrix and cell-cell communication that is essential for both epithelial and stroma repair/regeneration. Despite these positive outcomes for short term treatment, IL1RA or CMC did not restore collagen fiber alignment in the stroma at 24 hr, contrasting to the extensive restoration at 14 days of PBS treatment (Supplementary, Figure S5B), implying that remodeling of the collagen matrix requires an extended period of time.

Taken together, our results show lubrication alone is sufficient to restore the corneal stroma and BM during chronic inflammation in part through dampening IL1-MAPK signaling (Figure 7). Dampening keratocyte inflammation appears to be the first step in initiating a reparative process that includes the re-establishment of BM-ECM interactions and reorganization of the collagen matrix. Thus, our results highlight a key, and previously unappreciated role for topical lubrication as a therapeutic modality that rescues epithelial-matrix interplay and the associated network that supports both epithelial and stroma regeneration.

**Figure 7.**
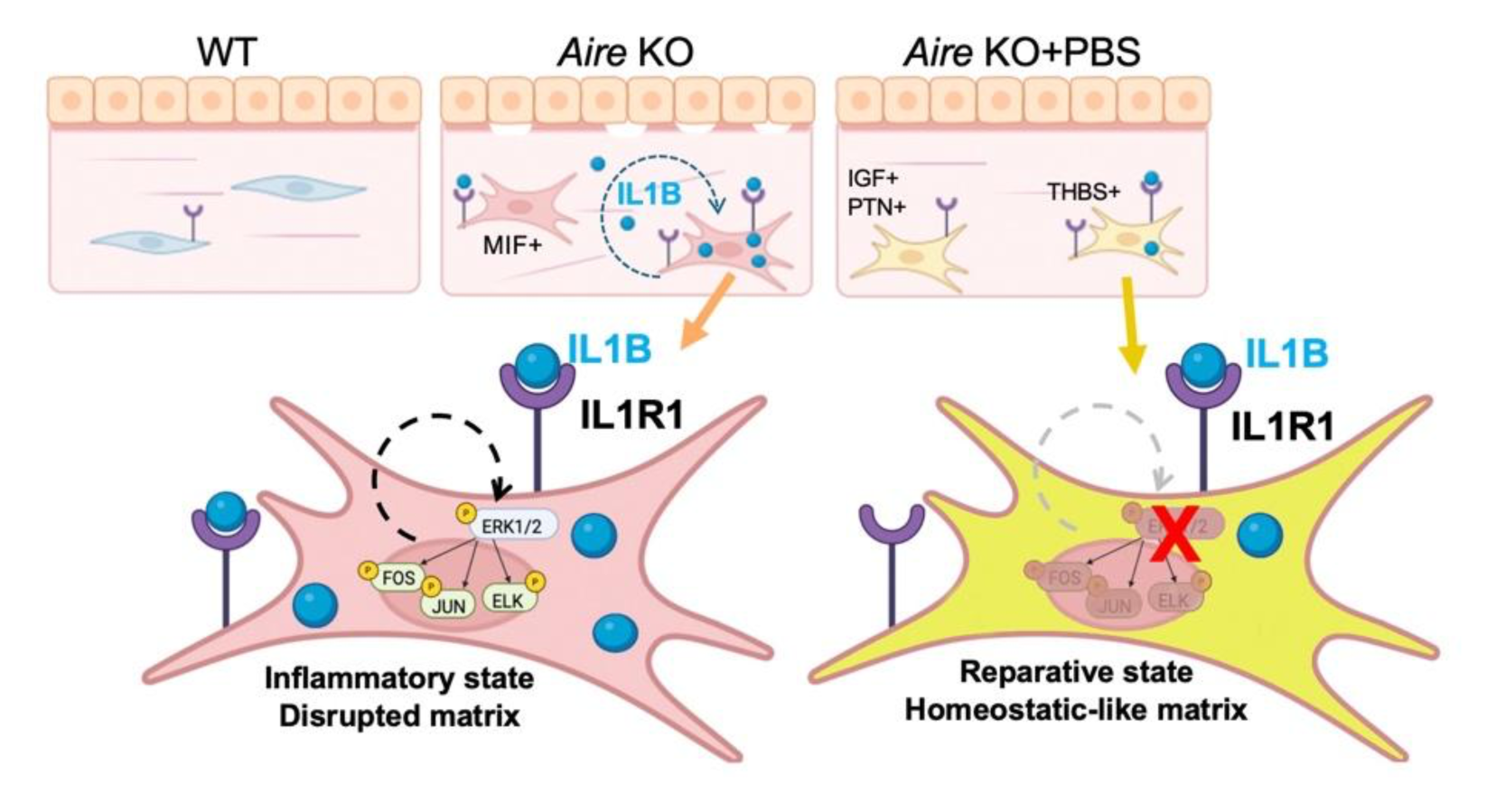
Lubrication promotes the transition of keratocytes from an inflammatory cell state to a reparative phenotype by suppressing the autocrine IL-1 loop activation. In the inflamed cornea, IL1β in the stroma triggers MAPK signaling in keratocytes and activation of the inflammatory/pro-fibrotic transcription factors FOS, JUN and ELK leading to fibroblast activation and matrix disruption. Topical PBS is sufficient to break this autocrine loop to generate keratocytes with a reparative phenotype.

## Discussion

Over-the-counter (OTC) topical lubricants are the most commonly prescribed treatment for symptoms of ocular dryness, but little is known about their therapeutic value for managing patients with dry eye disease. Here, we uncovered a previously unappreciated restorative function of topical lubrication in maintaining the stromal compartment of the desiccated cornea. Specifically, we show that application of the simplest form of eye drops (PBS) promotes the recovery of two matrices essential to corneal function, the BM and stromal ECM, resulting in a dramatic rescue of the precisely arranged collagen fibril network that is disrupted in aqueous-deficient dry eye.

Using transcriptomics, we discovered an unexpected level of stromal cell plasticity in response to prolonged desiccating stress and revealed the transition of distressed keratocytes from a destructive to reparative state with consistent and prolonged topical lubrication. We further discovered that lubrication achieves this outcome, in part, by dampening the well-established and powerful, proinflammatory IL1R1-pMAPK signaling pathway. Thus, we have identified an essential role for topical lubrication in maintaining the integrity of stromal compartment, as well as a key mechanism whereby tissue damage resulting from chronic desiccating stress can be reversed through inhibition of the IL-1/MAPK signaling cascade in stromal keratocytes. Upon injury, keratocytes are known to alter their phenotype giving rise to a population of cells that undergo rapid apoptosis or transition to an activated fibroblast-like state that is intended to prevent excessive inflammation and corneal opacification (8, 67). The transition to an inflammatory phenotype occurs in numerous inflammatory diseases such as rheumatoid arthritis, inflammatory bowel disease (IBD), Sjogren’s disease and ulcerative colitis (UC) (68, 69). Interestingly, this reprogramming is proposed as a two-step process, where exogenous inflammatory cytokines produced by immune cells trigger the transition of fibroblasts towards an inflammatory state which, in turn, produce high levels of inflammatory cytokines to amplify and sustain inflammation (68). Consistent with this process, our study identifies a subset of inflammatory keratocytes in the desiccated cornea that produce high levels of IL1b and co-express IL1R1, suggesting the potential activation of an autonomous inflammatory amplification loop. Activation of autocrine IL-1 production via an “IL-1 loop” has also been shown in corneal fibroblasts grown in vitro (70), further supporting a mechanism whereby activation of the IL-1/MAPK pathway in response to desiccating stress drives a sustained proinflammatory autocrine loop that induces the transition of these cells to an inflammatory remodeling phenotype. A similar mechanism has been reported in rheumatoid arthritis where a subset of synovial fibroblasts produces high levels of IL-6 (71) through an autocrine loop involving leukocyte inhibitory factor (LIF) and its receptor, LIFR, resulting in the continuous production of inflammatory mediators that impair disease resolution (72). Together, these studies suggest that targeting fibroblast amplification loops may be a novel and effective approach to attenuate fibroblast inflammatory responses and downstream tissue damage. Our data further suggest that consistent and ongoing lubrication of the diseased ocular surface is sufficient to break the inflammatory amplification loop in keratocytes thereby promoting the reprogramming of inflammatory keratocytes towards the reparative state.

To further unravel the regulatory pathway orchestrating keratocyte cell state transition in chronic dry eye disease we identified a set of pro-inflammatory/pro-fibrotic TFs that are highly enriched in these keratocytes. While some of these regulatory factors (e.g., NFKB, STAT, and IFN) have been reported as shared TFs enriched in inflammatory fibroblasts across a diverse group of immune disorders, others displayed distinctive TFs (e.g., FOSL2, JUN, and ELK4) unique to inflammatory keratocytes. Increased transcriptional activity of FOS, JUN and ELK have been previously linked to activation of IL1b signaling, further implicating this pathway as a key driver of keratocyte inflammation in dry eye. In contrast, we identified a set of pro-repair TFs enriched in keratocytes of the lubricated cornea. Several of these (e.g.,KLF9, NR2F2, and NFIA) are considered to be anti-inflammatory with KLF9 and NR2F2 specifically shown to inhibit fibroblast activation and inflammation (73). Other TFs, (e.g., FOXP1, EBF1, and NFATC1) are regulators of stem cell self-renewal and maintenance (74–76). Notably, EBF1 has been identified as a novel regulator of mesenchymal stromal progenitor cell maintenance and differentiation while NFIX and SOX4 promote wound repair and regeneration in muscle and skin, respectively (75, 77, 78). Interestingly, these specific pro-repair TFs were identified in a recent single cell sequencing study of dermal fibroblasts in a mouse model for wound-induced regeneration (49), further indicating that the lubricated dry eye keratocytes resemble regenerative fibroblasts.

Also unique to this investigation was our ability to pinpoint the specific spatial location of keratocyte clusters across the cornea under homeostatic and chronic inflammatory conditions. For example, the limbal niche is the location where limbal stem cells reside while also embodying the unique intersection of sensory nerves, blood vessels, and immune cells entering the cornea. We identified the presence of peripheral limbal-restricted keratocytes characterized by expression of *Fmod, Mmp3, Cxcl9*, or *Namp*t. Given their proximity to other critical players within the niche, it is likely they respond to signals from other niche cells (cytokines, growth factors, nutrients and neurotransmitters) which, in turn, transmit signals to other keratocytes within the corneal stroma. Keratocytes are postulated to form a functional communication network of interconnected cells across the cornea. These networks support the creation of centripetal signal transduction pathways whereby limbal keratocytes initiate cascades that activate central cornea keratocytes either to repair or disrupt stromal integrity. Hence, the limbal niche is critical for maintenance and renewal of the cornea and understanding the interactions between limbal keratocytes and the rest of the limbal niche will aid in developing targeted treatments for corneal diseases and improving clinical outcomes for patients with chronic inflammatory ocular diseases.

Our study provides the first robust evidence of the therapeutic benefit of topical lubrication in preserving and restoring ocular health. As the mainstay of dry eye therapies, we provide clear evidence that corneal lubrication is an essential part of dry eye management, thus providing much-needed guidance to practicing clinicians and patients. Our data also reveal for the first time a potential role for fibroblast-targeted therapies as a novel approach to restoring ocular surface health in dry eye. This is significant in that currently available FDA-approved anti-inflammatory agents are not sufficient to control the signs and symptoms of dry eye in the majority of patients with moderate to severe cases. Thus, the restorative properties of lubrication in combination with anti-inflammatory and fibroblast modulating therapies could revolutionize our future approach to dry eye management.

## Material and methods

### Resource availability

#### Lead contact

Further information and requests for resources and reagents should be directed to and will be fulfilled by the lead contact, Sarah Knox **Materials availability**

This study did not generate new unique reagents.

#### Data and code availability

RNA sequencing data have been deposited at the Gene Expression Omnibus (GEO) with accession code GSE272702 listed in the key resources table and are publicly available as of the date of publication.

Original microscopy data reported in this paper will be shared by the lead contact upon request.

This paper does not report original code.

Any additional information required to reanalyze the data reported in this paper is available from the lead contact upon request.

## EXPERIMENTAL MODEL AND SUBJECT DETAILS

### Animals

All procedures were approved by the UCSF Institutional Animal Care and Use Committee (IACUC) and adhered to the NIH Guide for the Care and Use of Laboratory Animals (Approval number: AN203203-00). Wild type (WT) and *Aire*-deficient mice on the BALB/c background (BALB/c *Aire* KO) were the gift of Mark Anderson, University of California, San Francisco. Adult female mice (aged 5-7 weeks) were used in all experiments. Mice were housed in groups of up to five per cage where possible, in individually ventilated cages (IVCs), with fresh water, regular cleaning, and environmental enrichment. Appropriate sample size was calculated using power calculations. Genomic DNA isolated from tail clippings was genotyped for the *Aire* mutations by PCR with the recommended specific primers and their optimized PCR protocols (Jackson Laboratories Protocol 17936).

## METHOD DETAILS

### Treatment Regimen

*Aire KO* female mice aged 5 weeks were used for the study. A set of *Aire* KO mice were topically treated with 5 μL per eye of PBS three times daily for 14 consecutive days. Corneas were harvested after 24hr of treatment and processed for global RNAseq, and after 14 days of treatment corneas were collected for *in-situ* and immunofluorescence analysis, and snRNAseq. For IL1R1 signaling inhibition study, 7wk Aire KO mice were topically treated with 5 µL per eye of 50 µg/mL Concentration IL1R1 antagonist (IL1RA) or vehicle control Carboxymethylcellulose (CMC) three times over 24 hours.

### Tissue processing and immunohistological analyses

#### *In situ* Hybridization

RNAscope^TM^ HiPlex Assay v2 (ACDBio) was performed using company protocols on fresh frozen cornea tissue sections to detect target RNA at single cell level. Tissue pre-treatment included fixation for 60 min in 4% paraformaldehyde (PFA) at RT, Dehydrate the sections in 50%, 70%, and 100% ethanol followed by protease IV treatment (RNAscope® Protease IV ACD# 322336) for 30 min at RT. Target probes were hybridized for 2 hrs at 40°C using the HybEZ Oven followed by amplification steps according to the manufacturer’s protocol. Briefly, treatment with ACD’s Amp 1 (Cat. 324111), Amp 2 (Cat. 324112), and Amp 3 (Cat. 324113) were performed for 30 min each at 40℃. Mouse Positive Control Probe (ACD Probe: 320881) and Negative Control Probe (ACD Probe: 320871) were included in the assay. Detection of specific probe binding sites was with RNAscope HiPlex12 Detection Reagents (488, 550, 650) kit v2 (ACDBio, Cat. 324410). Detection of the first 3 probes were with Fluoro T1-T3 (Cat. 324411) for 15 min at 40℃ and counterstained with DAPI (Cat. 324420). Positive staining was indicated by fluorescent dots in the cell cytoplasm or nucleus. The slides were mounted and imaged at 40x magnification on Zeiss Yokogawa Spinning disk confocal microscope. Fluorophores T4-T6 (Cat. 324412), T7-T9 (Cat. 324413), and T10-T12 (Cat. 324414) were added in subsequent rounds to the tissue. To remove the coverslips, the slides were soaked in 4X SSC and the fluorophore was removed using a 10% cleaving solution (Cat. 324130) incubated for 15 min at RT twice.

### Immunofluorescence analysis

Immunofluorescent analyses of cornea samples were performed as previously described (9). Briefly, enucleated eyes were embedded in OCT Tissue Tek freezing media. 7μm and 20 μm sections were prepared from fresh frozen tissues using a cryostat (Leica, lzar, Germany) and mounted on SuperFrost Plus slides. Sections were fixed for 20 min in 4% paraformaldehyde (PFA) at room temperature (RT) and permeabilized using 0.3% Triton X100 in phosphate buffered saline for 15 min. Sections were then washed in PBS-Tween 20 (PBST) for 10 min, before being blocked with 5% normal donkey serum (Jackson Laboratories, ME) in PBST for 1 hour at RT. After blocking, slides were incubated with primary antibodies diluted in blocking buffer overnight at 4°C. Following 3 washes with PBST, slides were incubated with secondary antibodies diluted in blocking solution at RT for 1 hour.

For immunofluorescent analysis, 7 or 20 μm tissue sections were incubated with the following primary antibodies: Rabbit anti-LAMA3 (1:500, Abcam, Cat 151715), rat anti-Perlecan (HSPG2) (1:500 Chemicon, MAB1948), rabbit anti-COL1(1:500, Abcam, Cat 34710), rabbit anti-CD34 (1:500, Abcam, Cat. ab81289), *mouse anti-pMAK (1:200, Upstate, Cat. 05-481), rabbit anti-pFAK (1:500, Abcam, Cat.223529), rabbit anti-IL1R1 (1:400, Santa Cruz, Cat. sc-689), goat anti-IL1b (1:20, R&D,* Cat. AF-401_NA). Antibodies were detected using Cy2-, Cy3- or Cy5-conjugated secondary Fab fragment antibodies (Jackson Laboratories), and nuclei were stained with Hoechst 33342 (1:3000, Sigma-Aldrich). Fluorescence was analyzed using a Zeiss LSM 900 confocal microscope or Zeiss Yokogawa Spinning disk confocal microscope with images assessed using NIH ImageJ software (79), as described below.

### Image Analysis

#### Basement membrane and extracellular matrix (ECM) analysis

Laminin (LAM) and perlecan (HSPG2) intensities were quantified on 300 µm sections of central corneal basement membrane ROI applying Tsai’s thresholding method (Moments)(ref), integrated densities within the ROI of the thresholded image were recorded. Alignment of collagen I fibers and fiber thickness were measured using CurveAlign CT-FIRE mode according to software protocol (80) and the angle of each extracted fiber was calculated and recorded by the software. Frequency distribution histograms of the different groups were created using PRISM (n>=5) and subjected to gaussian curve fit (Supplementary Fig 1A).

### Quantification of mechanosensors pFAK

pFAK fluorescent intensity was quantified on a ROI containing a 350 μm section of central cornea epithelial BM. Tsai’s thresholding method (Moments) was applied to the ROI of each cornea, and integrated densities within the ROI of the thresholded image were recorded.

### Quantification of keratocyte inflammation pMAK and IL1b

pMAK activity level was quantified on a ROI containing the stromal area under 350 μm section of central cornea epithelial BM. Tsai’s thresholding method (Moments) was applied to the ROI of each cornea, and percent area of threshold signal within the ROI of the thresholded image were recorded. IL1b high keratocytes were identified visually and manually counted within 1000um of cornea cross-section. To exclude immune cells that also expressed IL1b, only IL1b^high^IL1R1^+^ cells were counted because we have previously shown that immune cells in the cornea did not express IL1R1.

### Quantification of corneal stromal thickness

Stromal thickness was measured at the center of the cornea (the thickest region of the whole cornea) between corneal epithelial basement membrane and cornea endothelium, from 40x cornea cross section images.

### Quantification of the RNAScope Hiplex images

Image analysis was performed utilizing Qupath, a bioimage analysis software. Images taken at 40x magnification were uploaded to the software and regions of interest were drawn to include sections of the corneal stroma at limbus and central cornea. ROIs were 200um of in length for limbus, 350um in length for central cornea, and 500um for limbal peripheral cornea, and included the entire area between the corneal endothelium and epithelial basement membrane. Qupath’s cell detection feature was used to quantify nuclei staining and subcellular detection was used to quantify the number of RNA puncta within the cell. The threshold intensity for nuclei detection was set to 10, and the cell expansion was set at 5um from the detected nuclei. The thresholds for subcellular spot detection were determined using negative control images. Threshold values that yielded 10% or less of the total detections on the negative control (i.e. nuclei and puncta) were applied to the target images.

### Collagen Fiber alignment and thickness analysis

Images of the corneal stroma with Collagen I staining were analyzed using the Curvelet-Transform based fibrillar collagen quantification tool (CurveAlign and CT-FIRE) (80). The output measurement includes fiber angle, alignment, and thickness.

### RNA isolation and RNAseq library generation

Total RNA was collected at 7 days of treatment and purified using RNAaqueous and DNase reagents according to the manufacturer’s instructions (Ambion, Houston, TX, USA). RNA quality was assessed using the Agilent 2100 BioAnalyzer, and samples with an RNA integrity ≥ 6.0 were included for RNA sequencing. The synthesis of mRNA libraries was performed by Novogene Corporation Inc. according to their protocols. RNA library was formed by ployA capture (or rRNA removal), RNA fragmentation by covaris or enzyme digestion and reverse transcription of cDNA. Sequencing was performed as described below

### Bulk-RNAseq analysis

RNA libraries were sequenced on an Illumina NovaSeq 6000 by Novogene Corporation Inc. Depths of 20–30 million 150 bp paired-end reads were generated for each sample. Quality control metrics were performed on raw sequencing reads using fastp (81). In this step, clean data (clean reads) were obtained by removing reads containing adapter and poly-N sequences and reads with low quality from raw data. At the same time, Q20, Q30 and GC content of the clean data were calculated. All the downstream analyses were based on clean data with high quality. Reference genome and gene model annotation files were downloaded from genome website browser (NCBI/UCSC/Ensembl) directly. Paired-end clean reads were aligned to the reference genome using the Spliced Transcripts Alignment to a Reference (STAR) software (82). DEseq2 was then used to detect differential gene expression between WT, untreated Aire KO, PBS treated Aire KO and Lacripep treated Aire KO corneas based on the normalized count data. Genes were considered differentially expressed if the log2 Fold Change between samples was at least 1, with the adjusted p-value held to 0.05 (83). Heatmaps and Volcano plot of differentially expressed genes were created using “pheatmap” (84), and “EnhancedVolcano” (85) R packages, respectively.

### Single nuclei (sn) RNA-sequencing

To ensure biological diversity, nuclei were initially isolated from snap frozen corneas of 5 mice per treatment group at 7 wks of age (WT, KO, PBS, LAC). Nuclei isolation protocol was adapted from 10x Genomics (Sample Preparation Demonstrated protocol, Rev D). Briefly, fresh corneas were dissected, minced, snap frozen in lysis buffer (Sucrose 0.32M, CaCl_2_ 3mM, Mg(Ac)_2_ 3mM, EDTA 0.1mM, Tris-HCL 10mM, DTT 1mM, Triton x100 0.1%, DEPC-treated water) and stored for further processing. Upon nuclei isolation, frozen sample were crushed using a pastel and combined into 15ml tube, with total volume of 5ml lysis buffer. Samples kept on ice for 15min with “vortexing” every 5 min. Sample were then transferred into ultracentrifuge tubes on ice (Beckman Coulter 355631), and 9ml of Sucrose solution (Sucrose 1.8M, Mg(Ac)_2_ 3mM, Tris-HCL 10mM, DTT 1mM, DEPC-treated water) were carefully added to the bottom of the tube. Sample were spined 24,400 rpm for 2.5hr at 4°c. After spin, supernatant was removed, and 200ml of DEPC-based PBS was added for 20 min incubation on ice followed by resuspension. isolated nuclei solution was filtered twice using 10mm filter. Nuclei were counted before subjection to single-cell RNA sequencing using the Chromium Single Cell 3’ Reagent Version 3 Kit (10× Genomics). Sequencing was performed on Illumina HiSeq 2500 according to the 10× Genomics V3 manual.

### Single Nuclei RNA-seq analysis

Reads were aligned to the Mouse reference genome (mm10-2020-A) using STAR and resulting bam files were processed with the Cell Ranger pipeline v6.1.2, followed by removal of ambient RNA contamination using DecontX. DecontX, is a Bayesian method, which deconvolutes the expression matrix into a matrix of native counts for each cell, and a matrix of contaminated counts, originating from other cell populations. Using Seurat v4.0 in Rstudio we filtered out low-quality cells based on UMI counts and the number of genes. The batch correction was performed with Seurat v4.0. Clustering was performed with CellFindR v2.0.0 using settings with a quality measure of ≥5 genes with FC>1.5. Subsequent analysis was performed with Seurat v4.0. Clusters were visualized using Uniform Manifold Approximation and Projection (UMAP) dimensional reduction (RunUMAP). Markers for each cluster were identified with FindAllMarkers using default parameters. Stroma clusters were identified using the known markers Dcn and Lum and extracted from the entire cornea dataset for further analysis using Seurat and CellFindR. Regulon activity was assessed with SCENIC v1.1.2.4, SCENIC combines GENIE3 (86) with regulatory binding motif enrichment to simultaneously cluster cells and infer regulatory networks. For cell-cell ligand-receptor interaction analysis, Seurat objects of corneal stroma data were analyzed using CellChat21 using standard parameters. Cytoscape 3.9.1 was used to visualize TF-target networks.

### Quantification and Statistical Analysis

All data are expressed as mean + SD. A minimum of three independent repeats were conducted in all experiments. Bar graphs were used to summarize the means and standard deviations of each outcome obtained using all data collected from wild type (WT) and *Aire KO* mice. A Student’s t-test was used for two group comparisons; a one-way analysis of variance (ANOVA) was used for multiple group comparisons with either Tukey’s multiple comparisons test, Dunnett T3, or corrected for multiple comparisons by controlling the False Discovery Rate using Benjamini, Krieger and Yekutieli; P < 0.05 was considered statistically significant. A false discovery rate of 0.05 was applied to bulk-RNAseq data. A Wald test with Benjamini and Hochberg correction was used for differential gene expression.

## Acknowledgements

The authors would like to thank Bo Sun for helping to process our snRNAseq dataset on HPC cluster. The authors would also like to thank Drs. Suzanne M.J. Fleiszig (University of California, Berkeley), and Matilda Chan (University of California, San Francisco), and our undergraduate student volunteer Darleene Nguyen for their constructive input toward this manuscript. This work was supported by the following funding sources: National Eye Institute grants R01EY025980 (to S.M.K. and N.A.M.), R01EY027392 (to S.M.K.), and R01EY033040 (to S.M.K. and N.A.M.).

## Author Contributions

Conceptualization, Y.E., F.Y.T.C., N.A.M., and S.M.K.; methodology, Y.E., F.Y.T.C., K.N.C., D.F, N.A.M., and S.M.K.; investigation, Y.E., F.Y.T.C., K.N.C., and D.F.; data acquisition, Y.E., F.Y.T.C., K.N.C., D.F., V.N., H,S., and L.A.; visualization, Y.E., F.Y.T.C., and S.M.K.; funding acquisition, S.M.K. and N.A.M.; project administration, S.M.K. and N.A.M.; supervision, S.M.K. and N.A.M.; writing – original draft, Y.E., F.Y.T.C., N.A.M., and S.M.K.; writing – review & editing, Y.E., F.Y.T.C., N.A.M., and S.M.K.

## Declaration of Interests

All authors declare no competing interests.

---

**Supplementary Figure S1.**
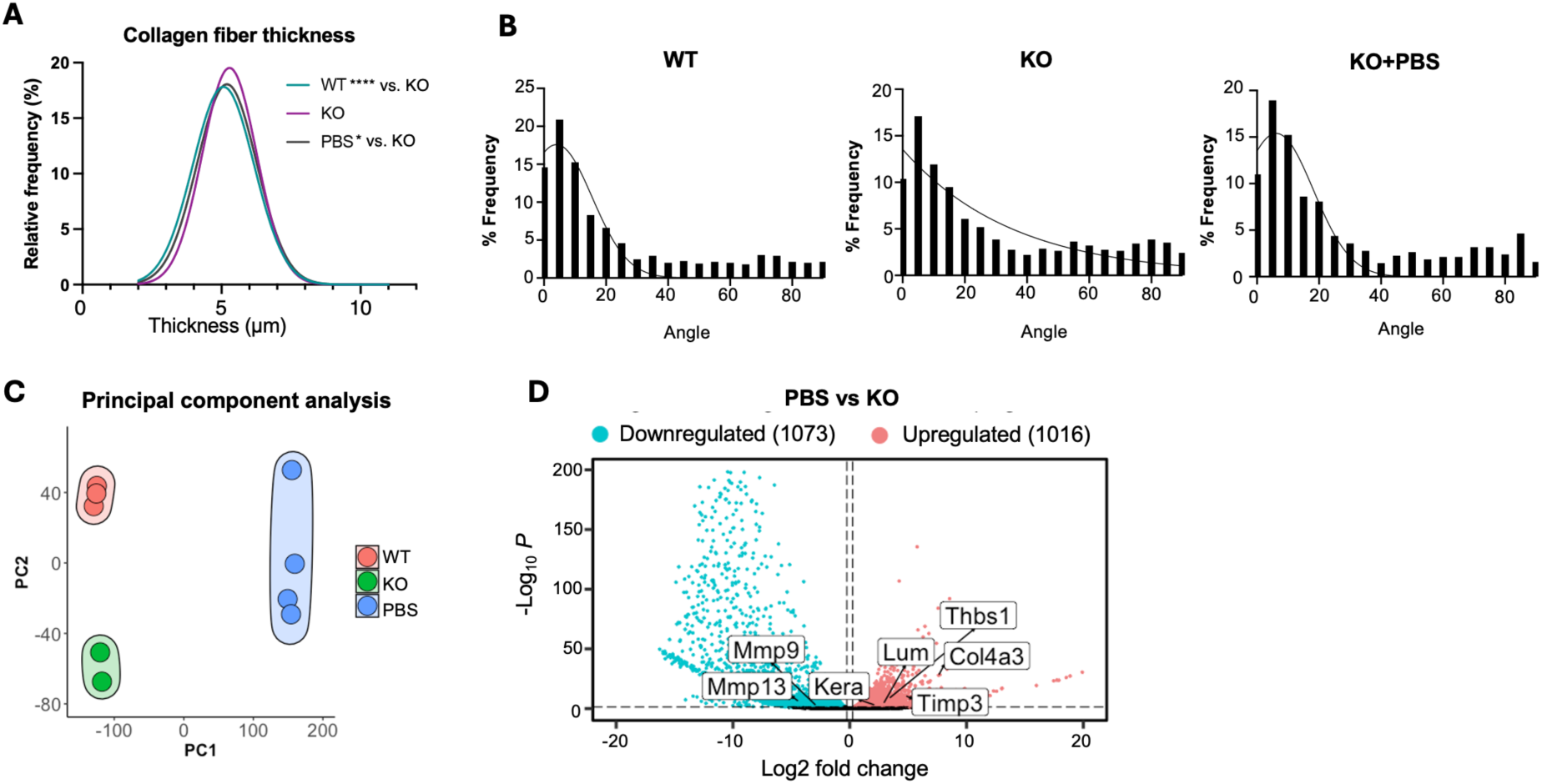
Topical lubrication of the ocular surface restores the basement membrane and promotes stromal regeneration, related to Figure 1. **A-B.** Distribution of the collagen 1 fiber thickness (**A**) and alignment angle in relation to the BM (**B**) in the corneal stroma of each group. **C.** Principal component analysis (PCA) of the different treatment groups. **D.** Volcano plot of differentially expressed corneal genes in response to PBS versus untreated KO after 24hr of treatment. Red and blue dots represent significantly upregulated and downregulated genes, respectively (log2 FC ≥1 and padj <0.05). Data in B were subjected to a Kolmogotrov-Smirnov test. Each dot in the bar graph represents a biological replicate. n > 4 mice per group except for 24hr RNAseq analysis where n ≥ 2.

**Supplementary Figure S2.**
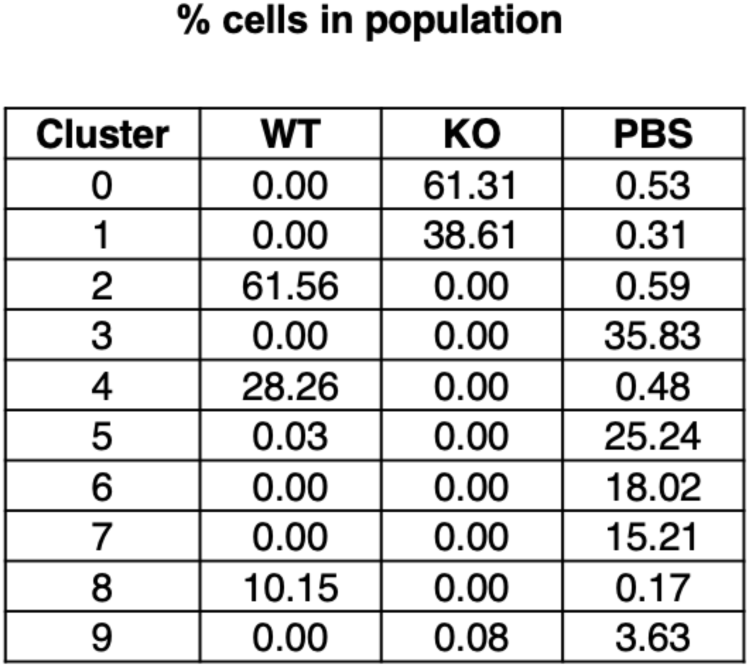
Table summarizing the proportion of keratocytes in each cluster within the different groups, related to Figure 2.

**Supplementary Figure S3.**
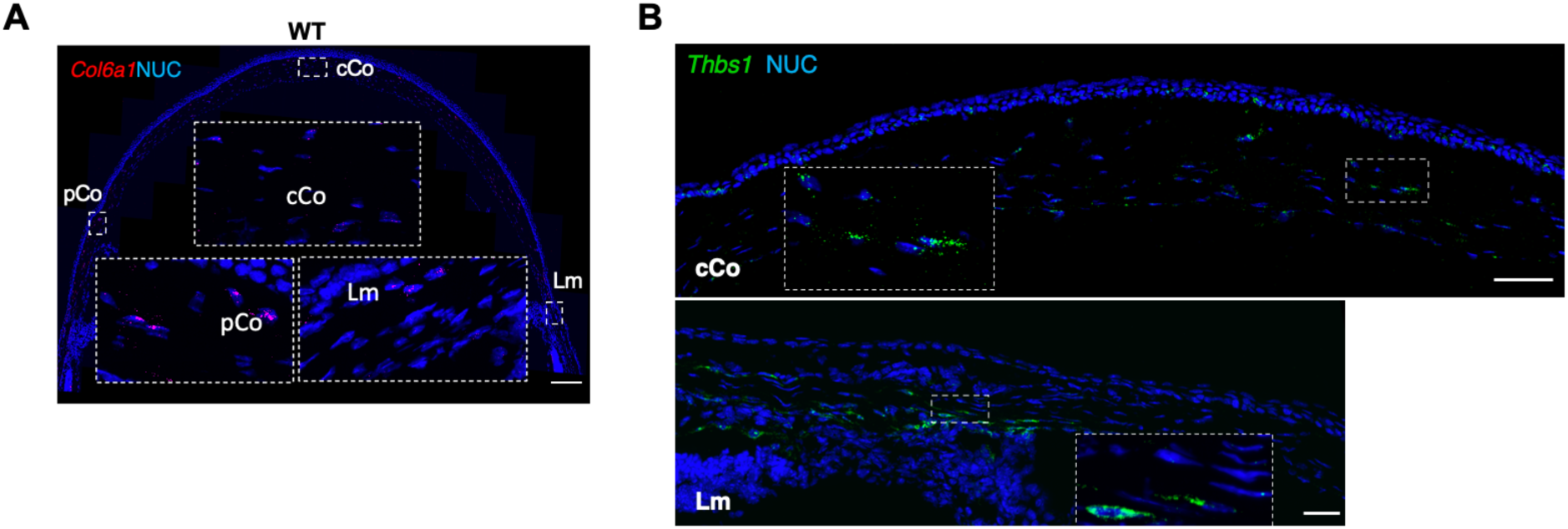
Spatial locations of *Col6a1+* and *Thbs1+* cells in the central, peripheral and limbal regions of the wild type (WT) cornea, related to Figure 3. Scale bars = 50 μm.

**Supplementary Figure S4.**
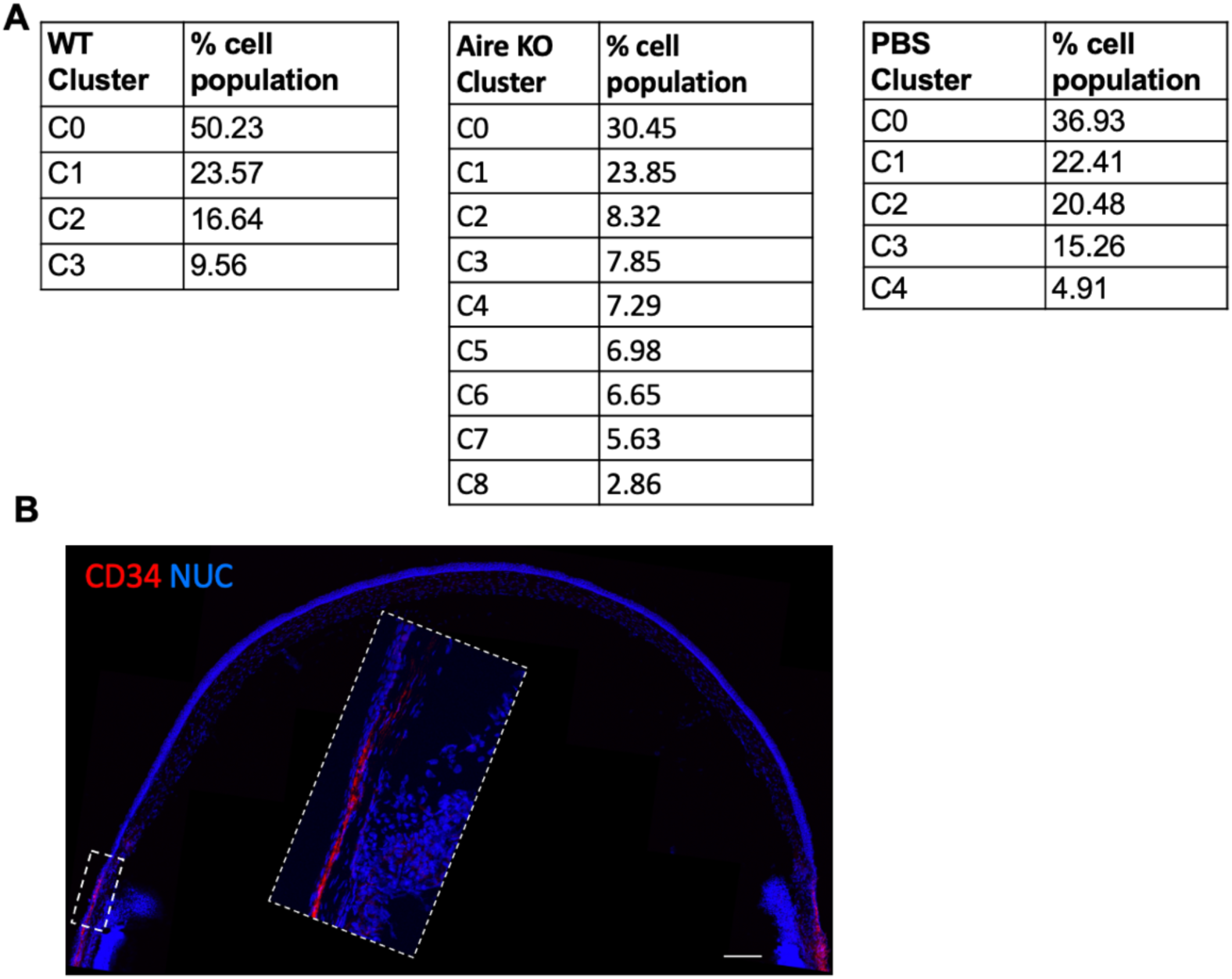
Table summarizing the proportion of keratocytes in each cluster within the different groups identified by snRNAseq (**A**) and the location of CD34+ cells at the limbal region (**B**), related to Figure 4.

**Supplementary Figure S5.**
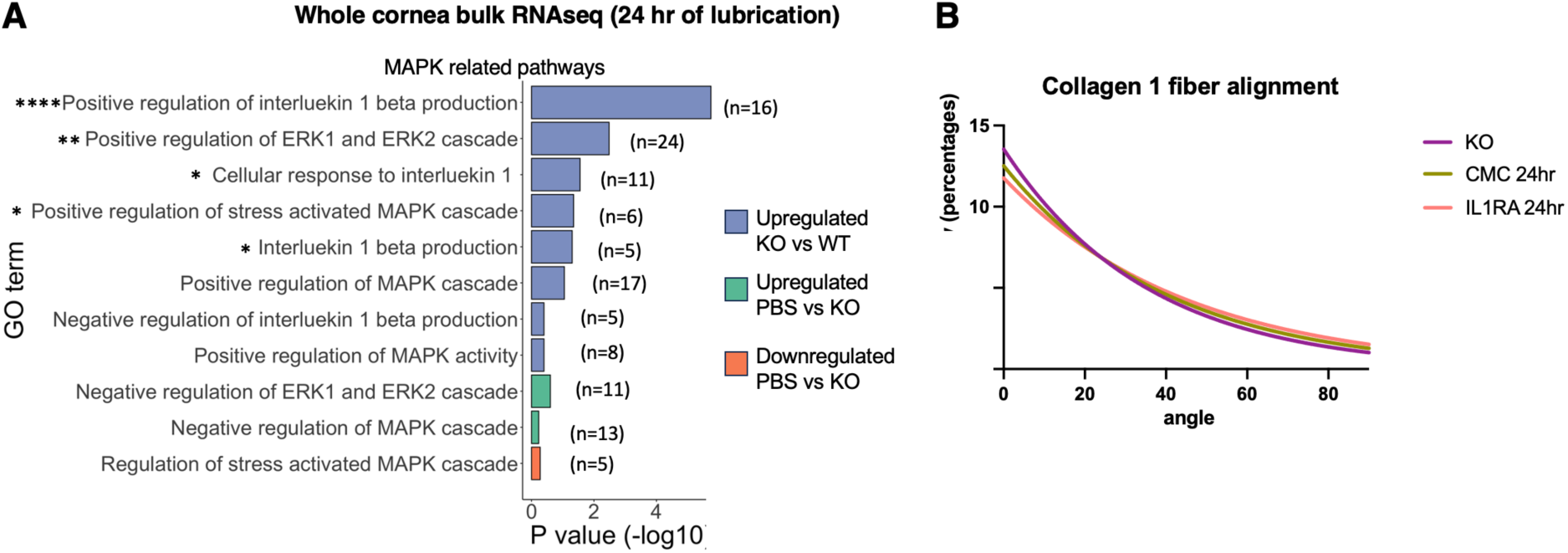
Lubrication rescues stromal architecture via inhibition of IL1B-IL1R1-MAPK signaling, related to Figure 6. **A.** Gene ontology (GO) analysis highlighting alterations in IL1 and MAPK related pathways in WT, KO and KO+PBS stromal samples at 24 hr of treatment (bulk RNAseq). **B.** Frequency of collagen 1 fiber alignment, comparing KO, KO+CMC and KO+IL1RA at 24hr.

## Notes

### Competing Interest Statement

The authors have declared no competing interest.

